# Natural age-related sleep-wake alterations onset prematurely in the Tg2576 mouse model of Alzheimer’s disease

**DOI:** 10.1101/2021.10.25.465747

**Authors:** Sedef Kollarik, Carlos G. Moreira, Inês Dias, Dorita Bimbiryte, Djordje Miladinovic, Joachim M. Buhmann, Christian R. Baumann, Daniela Noain

## Abstract

Sleep insufficiency or decreased quality have been associated with Alzheimer’s disease (AD) already in its preclinical stages. Whether such traits are also present in rodent models of the disease has been poorly addressed, somewhat disabling the preclinical exploration of sleep-based therapeutic interventions for AD. We investigated age-dependent sleep-wake phenotype of a widely used mouse model of AD, the Tg2576 line. We implanted electroencephalography/ electromyography headpieces into 6 months old (plaque-free, n=10) and 11 months old (moderate plaque-burdened, n=10) Tg2576 and age-matched wild-type (WT) mice and recorded vigilance states for 24 hours. Tg2576 mice exhibited significantly increased wakefulness and decreased non-rapid eye movement sleep over a 24-hour period compared to WT mice at 6, but not at 11 months of age. Concomitantly, delta power appeared decreased in 6-month old Tg2576 mice in comparison to age-matched WT controls, yielding a reduced slow-wave energy phenotype in the young mutants. Lack of genotype-related differences over 24 hours in overall sleep-wake phenotype at 11 months of age appears to be the result of the natural aging of WT mice. Therefore, our results indicate that at plaque-free stages of the disease, diminished sleep quality is present in Tg2576 mice which resembles aged healthy controls, suggesting an early-onset of sleep-wake deterioration in murine AD. Whether such disturbances in the natural patterns of sleep could in turn worsen disease progression warrants further exploration.

## 1. Introduction

Altered sleep-wake patterns and reduced quality of sleep are associated with Alzheimer’s disease (AD), the most common neurodegenerative disease worldwide, already in its preclinical stages. Sleep-wake disturbances in AD include circadian shifts in sleep-wake cycle, excessive daytime sleepiness, significant decrease in time spent in both non-rapid eye movement (NREM) sleep - also known as slow-wave sleep (SWS) - and rapid-eye movement (REM) sleep, as well as increased sleep fragmentation.^1–3^

Human studies suggest that AD pathology targets circuits involved in normal sleep-wake behavior. For instance, degeneration of locus coeruleus, which produces norepinephrine and plays a crucial role in the sleep/wake switch, is already present in early stages of AD.^4, 5^ Likewise, functional impairments in the hippocampus, a brain area known to harbor sleep-mediated memory formation, were detected in healthy people at genetic risk for AD.^6, 7^ On the other hand, evidence also suggests that poor sleep is a risk factor for AD. Functional imaging and cerebrospinal fluid profiling in healthy elderly volunteers revealed that greater neurotoxic burden and higher amyloid pathology biomarker levels are associated with shorter sleep duration and poorer sleep quality, respectively.^8, 9^ This findings were further corroborated by a recent meta-analysis indicating that individuals with sleep disturbances had a ~4 times higher risk of preclinical AD than those without sleep complaints.^10^ Furthermore, a study in a murine transgenic AD model has shown that sleep deprivation increases the accumulation of amyloid plaques, while sleep induction reduces it,^11^ hinting sleep as a plausible target of interest in AD. Altogether, although sleep-wake disturbances have long been believed to be symptoms of AD, increasing evidence suggests a bidirectional relationship between sleep and AD. Overall, these findings emphasize the importance of studies focusing on sleep enhancement in at-risk populations, even before the neuropathological hallmarks of AD (i.e. amyloid plaques and tau tangles) set in.

Although human subjects are commonly used in studies focusing the relationship between sleep and neurodegeneration, animal models of AD undoubtedly provide relevant insights into the aforementioned associations between sleep-wake behavior and neurodegeneration in AD in ways that are not possible in human volunteers. Indeed, there are several studies examining the sleep-wake phenotype of AD mouse models. Zhang and colleagues (2005) observed a disruption of REM sleep related to plaque formation around the mesopontine cholinergic neurons in the Tg2576 line.^12^ In the same model, circadian period assessed by measuring wheel-running rhythms in constant dark was also found to be disrupted compared to the wild-type (WT) controls.^13^ In AβPP/PSEN1 mice compared to PSEN1 and WT mice, significant and progressive increase in the time spent awake, as well as an increase of theta power together with a decrease of delta frequencies was displayed by hippocampal electroencephalography (EEG).^14^ However, the highly spatially circumscribed expression of amyloid plaques in this model raises the question of whether such results can be extrapolated to other models of the disease and, more importantly, to AD patients. A more recent study examining three different mouse models (3xTgAD, Tg2576, and APP/PS1) demonstrated that both Tg2576 and APP/PS1 mice showed stage-dependent shifts in EEG power spectrum despite the time spent in each state being similar between the AD animals and their WT controls, while there were no power spectrum shifts in the 3xTgAD mouse line.^15^ However, the study did not focus on age-dependent changes therefore, disease progression as a potentially determinant factor on sleep-wake phenotype was not examined. Taken together, only a handful of studies have explored progressive sleep-wake alterations in mouse models of AD with no consensus regarding the nature of the relationship between sleep disturbances and disease progression (i.e. plaque-free vs. plaque-burdened stages). Moreover, the current knowledge on the effect of light-dark cycle on EEG-defined sleep-wake properties as well as other important EEG characteristics during SWS is rather scarce in murine AD.

In this study, we investigated the sleep-wake behavior of Tg2576 mice^16, 17^ via 24-h electroencephalography/ electromyography (EEG/EMG) recordings at two early stages of disease progression: plaque-free, pre-morbid stage, and moderately plaque-burdened disease stage and compared both groups to age-matched WT controls. We designed this experiment to provide insights into the specific changes in sleep-wake behavior and characteristics depending on i) circadian aspects (light vs dark periods), ii) genotype, and/or iii) age (i.e. stage of disease progression), in a well-characterized mouse model of AD.

## 2. Methods

### 2.1. Animals

We examined male and female Tg2576^16^ mice overexpressing a mutant form of amyloid precursor protein (APP), APPK670/671L, linked to the early-onset familial AD and their non-transgenic WT littermates (Taconic Biosciences; Cologne, Germany). We excluded the mice carrying the Pde6brd1 retinal degeneration allele, which may lead to retinal degeneration resulting in light sensitivity and/or blindness. The study design included both male and female mice at pre-morbid, i.e. plaque-free (6 months old Tg2576 mice, n = 10, and age-matched WT mice, n = 10), and moderate, i.e. plaque-burdened (11 months old Tg2576 mice, n = 10, and age-matched WT mice, n = 10) disease stages. We group-housed female mice, but kept males individually caged due to aggressive behavior. The animal room temperature was constant at 21 – 23 °C, with a 12:12 h light:dark cycle starting at 8.00 or 9.00 a.m., according to season. Mice had access to food and water *ad libitum* and had daily routine health checks throughout the study. All experiments were approved by the veterinary office of the Canton Zurich and conducted according to the local and federal guidelines for care and use of laboratory animals under license ZH210/17.

### 2.2. Surgery

We performed all surgical procedures under deep inhalation anesthesia with isoflurane (4 - 4.5 % for induction in anesthesia box, 2 - 2.5 % for maintenance using a nose cone fitting), and applied lidocaine, a local anesthetic (Xylocaine, Zürich) on the surface of the head skin prior to surgery. At the end of the surgery, we administered a combination of an anti-inflammatory (5 mg/kg, s.c., Metacam®, Zurich) and a pain relief drugs (0.1 mg/kg, s.c., Temgesic®, Zurich) to further prevent postoperative inflammation and pain.

The animals were implanted as described before.^18^ We placed 2 stainless steel screws (Bossard, #BN650, 1421611), one for each hemisphere, located 2 mm posterior to Bregma, 2 mm lateral from midline, connected to a pin header (Farnell, #M80-8530445) for EEG recording, and fixed the structure with dental cement (**Figure 1.A**.). We then inserted two gold wires bilaterally into the neck muscles for EMG recording and applied sutures to close the skin around the implant. Postsurgical analgesia was administered throughout the three days following the surgery during both light (0.1 mg/kg, s.c., Temgesic®, Zurich) and dark (1 mg/kg via drinking water, Temgesic®, Zurich) phases. We monitored body weight and home cage activity daily during the first week after the surgery and once per week thereafter.

**Figure 1.**
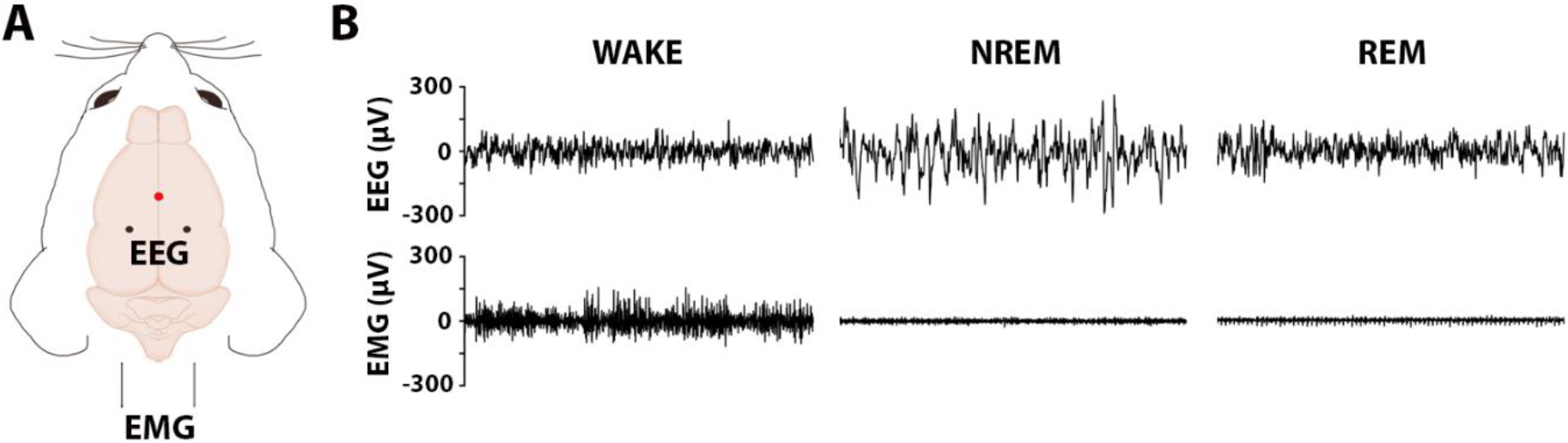
Illustration of electrodes positioning and raw EEG/EMG signals. **A)** Schematic of EEG/EMG implantation in a mouse (for details see Materials and Methods). Red circle (bregma) shows our reference point for the EEG electrode implantation and black circles reflect the EEG electrodes in both hemispheres. **B)** Representative EEG/EMG signal (2 epochs) in three vigilance states.

### 2.3. Data acquisition

To characterize sleep-wake behavior and EEG characteristics in freely moving Tg2576 and WT mice, 24 h EEG/EMG recordings took place at 6 months old (6-mo) mice, i.e. in a pre-morbid stage, and for assessments in moderate stage in 11 months old (11-mo) mice. Signals were amplified using an N700 polysomnography (PSG) amplifier (Embla, Ontario, Canada) with a bipolar montage, digitized and collected at a sampling rate of 200 Hz and an input range of ±200 mV using a 16-bit digital-to-analog-converter. We used Somnologica Science, version 3.3.1 (Resmed, Saint-Priest, France) for data collection.

### 2.4. EEG post-processing

We analyzed EEG/EMG recordings using SPINDLE (Sleep Phase Identification with Neural networks for Domain-invariant Learning).^19^ The software determines three vigilance stages **(Figure 1B)** in time slots of 4 seconds per epoch. Wakefulness (WAKE) proportions were calculated based on increased EMG activity for more than 50% of a 4-s epoch. NREM proportions were calculated by reduced EMG activity and increased EEG power in <4Hz frequency, while REM proportions were characterized by low EEG power in >4Hz oscillations and high EEG power in 6 - 9Hz frequency bands, with intermediate muscle tone. Transitions, such as wake to REM and REM to NREM were artificially suppressed. Following sleep scoring, we determined time spent, total bout number, mean bout duration and bout duration distribution in WAKE, NREM, and REM stages during the light-dark period for each mouse. Furthermore, we extracted measures of power in specific bandwidths, by processing the EEG signal on a custom MATLAB routine (ver. R2016b). Briefly, EEG signal was filtered between 0.5 – 30 Hz using a zero-phased Hamming-windowed linear FIR filter (order 100). Artifact data points within NREM epochs were removed. We performed a spectral analysis of consecutive 4 s epochs (FFT routine, Hamming window, 2 s overlap, and resolution of 0.25 Hz) and normalized the data indicating the percentage of each bin with reference to the total spectral power between 0.5 and 30 Hz.

### 2.5. Congo red staining and stereology

40 μm thick sagittal sections were mounted on slides and air-dried overnight. We stained the sections to identify the amyloid plaques in the brain tissue as previously described (Wilcock, Gordon, & Morgan, 2006). Briefly, on the day of staining, we rehydrated the sections by dipping the slides for 30 sec in distilled water. Then, we incubated the slides first for 20 min in the alkaline saturated NaCl solution (2 ml of 1 M NaOH solution into 200 ml of saturated NaCl was added previously to produce an alkaline solution), and then for 30 min in Congo red solution. Following that we quickly dehydrated the sections in 95% and 100% ethanol, cleared them in Roticlear®, and cover slipped with mounting reagent.

5 sagittal sections per mouse were used for stereological estimation of plaque burden in the hippocampus. A blinded experimenter counted the plaques with the optical fractionator technique using Stereo Investigator (MBF Bioscience) software. The plaques were visualized with Zeiss Imager M2 (63× oil immersion objective lens), the size of counting frame was 80 x 80 μm and the systematic random sampling grid size was 150 × 150 μm. The volume of the hippocampus (μm3) and the volume of each plaque (μm3) were assessed by the Stereo Investigator software.

### 2.6. Statistical analysis

Statistical analyses were conducted using IBM SPSS® software, and the figures were prepared in GraphPad Prism 8 (GraphPad Software, Inc., San Diego, CA). We detected the outliers by inspection of a boxplot for values greater than 1.5 box-lengths from the edge of the box, while normal distribution of variables was determined by Skewness and Kurtosis (p > .05). Levene’s test was used to assess the equality of variances.

For sleep proportion analysis, we conducted a two-way ANCOVA to examine the effects of genotype and age on sleep proportions, after controlling for sex. This was followed by pairwise comparisons. Due to abnormal distribution, we performed square root transformation for the REM sleep proportions.

Due to lack of normality, we ran a non-parametric Mann-Whitney U test to analyze sleep fragmentation. Specifically, we investigated the differences in the number of bouts and mean bout duration between genotypes in three vigilance states during light and dark periods of the day for each age category. We detected an outlier in REM bouts during dark period. Since the interpretation of the results was not impacted by the presence of the outlier, we decided to report the analysis including this animal. Subsequently, we examined the distribution of WAKE, NREM, and REM bouts in light and dark period, respectively, by dividing the bout durations into two categories: short bout durations (0-40 s) and long bout durations (>40 s). We calculated the relative number of bouts by dividing the number of bouts within these two categories by the total number of bouts for each mouse. Mixed-ANOVA was used to analyze the differences in bout distributions between the genotypes.

For power spectrum analysis, we conducted a two-way ANCOVA to examine the effects of genotype and age on preselected frequency bands: delta (0.5 – 4 Hz), theta (4 - 8 Hz), alpha (8 – 12 Hz), sigma (12 - 16 Hz), and beta (16 - 30 Hz) in three vigilance states, separately. The analysis was followed by pairwise comparisons. Delta power within the NREM epochs was summed over 12 h light or dark periods and the differences between the genotypes were investigated by a two-way ANCOVA.

To examine the differences in slow wave energy (SWE), which was calculated as a cumulative sum of delta power in NREM sleep across 24 h recording, we ran a two-way mixed ANOVA (genotype*time or age*time). Mauchly’s test of sphericity indicated that the assumption of sphericity was violated for the two-way interaction. Therefore, we interpreted the results by using Greenhouse-Geisser correction.

NREM delta power density (power/minute) was calculated by dividing total NREM delta power with the time spent in NREM (in minutes), in both light and dark periods, and then analyzed with a two-way ANCOVA, after controlling for sex.

Plaque burden in each mouse was calculated by dividing the volume of total plaques in the region of interest (hippocampus) by the total volume of the region of interest. Then we ran independent samples t-test to examine the effect of genotype on plaque burden in each age group separately.

## 3. Results

### 3.1. Increased wakefulness during dark period in Tg2576 mice

Sleep-wake disturbances are highly common in AD patients, and changes in sleep appear to lead to deterioration of cognitive symptoms.^20^ Therefore, our first question was whether there are alterations in sleep-wake proportions in Tg2576 compared to their WT controls. We analyzed 24-h EEG/EMG recordings for percentage of time spent in three vigilance states (WAKE, NREM, and REM) in the light and dark periods separately, as well as per 24 h.

We observed no significant differences between Tg2576 and WT mice, regardless of age, for the percentage of time spent in WAKE, NREM sleep and REM sleep during the light period (**Figure 2A-C**).

**Figure 2.**
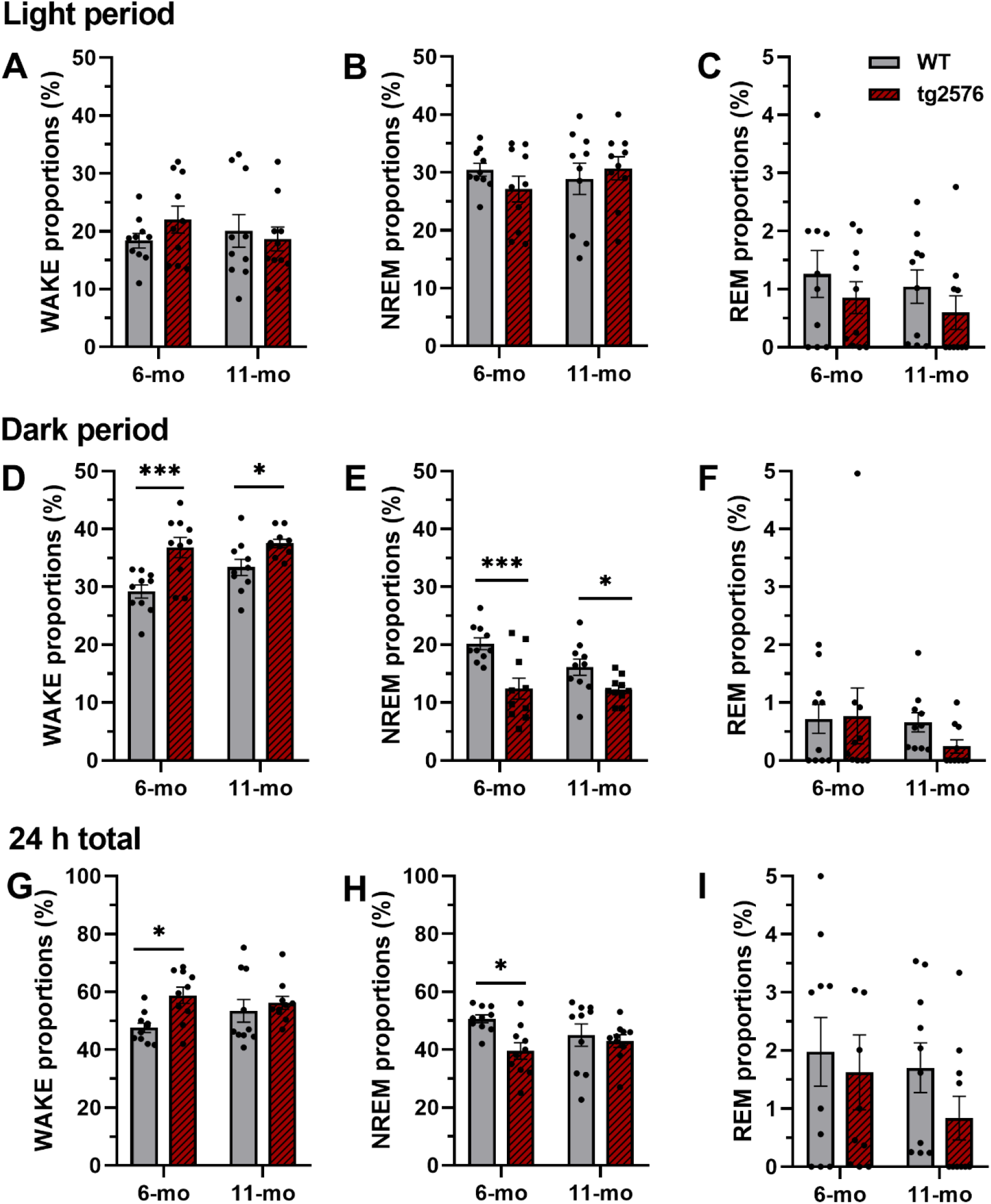
Sleep-wake proportions in Tg2576 mice compared to WT controls. There were no differences in **A)** WAKE, **B)** NREM, and **C)** REM sleep proportions between genotypes during the light period. In the dark period, Tg2576 mice showed increased **D)** WAKE and decreased **E)** NREM sleep, while **F)** REM sleep proportions were similar. Analysis of 24 h recordings showed an increase in **G)** WAKE and a decrease in **H)** NREM sleep only in 6-mo Tg2576 mice in comparison to WT controls. No difference was observed in **I)** total REM sleep proportions between genotypes. All data are expressed as mean ± SE-, \**p* < .05, \*\**p* < .01, \*\*\**p* < .001, two-way ANCOVA.

We observed, however, a statistically significant main effect of genotype in WAKE (*F*(1,35) = 20.54, *p* <.001, η_p_^2^ = .370) and NREM (*F*(1,35) = 20.42, *p* <.001, η_p_^2^ = .368) proportions during the dark period, after controlling for sex and when disregarding the effect of age. At 6 months, the gender-adjusted mean WAKE proportion for the dark period in Tg2576 mice was higher than in WT controls (95% CI [3.276, 10.588], *p* < .001, **Figure 2D**). The result persisted at 11 months as well, with higher WAKE proportions in Tg2576 (95% CI [0.823, 8.038], *p* = .018). Dark period NREM proportions in 6-mo and 11-mo Tg2576 mice were lower compared to age-matched WT controls (6-mo; 95% CI [3.301, 10.488], *p* < .001, 11-mo 95% CI [0.696, 7.788], *p* = .020, **Figure 2E**), whereas REM sleep was similar in both groups (**Figure 2F**).

The 24 h analysis revealed a higher WAKE and lower NREM proportion in 6-mo Tg2576 compared to WT controls (WAKE: 95% CI [1.333, 18.037], *p* = .024; NREM: 95% CI [1.109, 17.355], *p* = .027, **Figure 2G and H**), but not in 11-mo mice. REM sleep did not differ between genotypes in the 24 h period either (**Figure 2I**).

### 3.2. NREM sleep stage continuity appears to be impaired in Tg2576 mice

Since malfunction in sleep stability is considered just as hazardous to several aspects of health as short sleep duration,^21^ we further analyzed sleep fragmentation, a common sleep-wake disturbance associated with cognitive decline in AD patients,^22^ in Tg2576 mice. We determined the existence of sleep-wake instability by calculating the number of bouts and the mean bout duration within each vigilance state in both light and dark periods. Our results showed no significant differences in these measures between genotypes in both age groups (data not shown).

We further analyzed sleep stage continuity by focusing on the bout-length distributions in WAKE, NREM, and REM states. Our results indicated trend towards higher amount of short and lower amount of long bouts only in NREM in 6-mo Tg2576 mice both in light (*F*(1,18) = 4.260, *p* =.054, η_p_^2^ = .191) and dark (*F*(1,18) = 3.602, *p* = .074 η_p_^2^ = .167) periods when compared to age-matched WT littermates (**Figure 3A**). There were no differences in 11-mo mice (**Figure 3B**) in bout-length distributions of all three states between genotypes.

**Figure 3.**
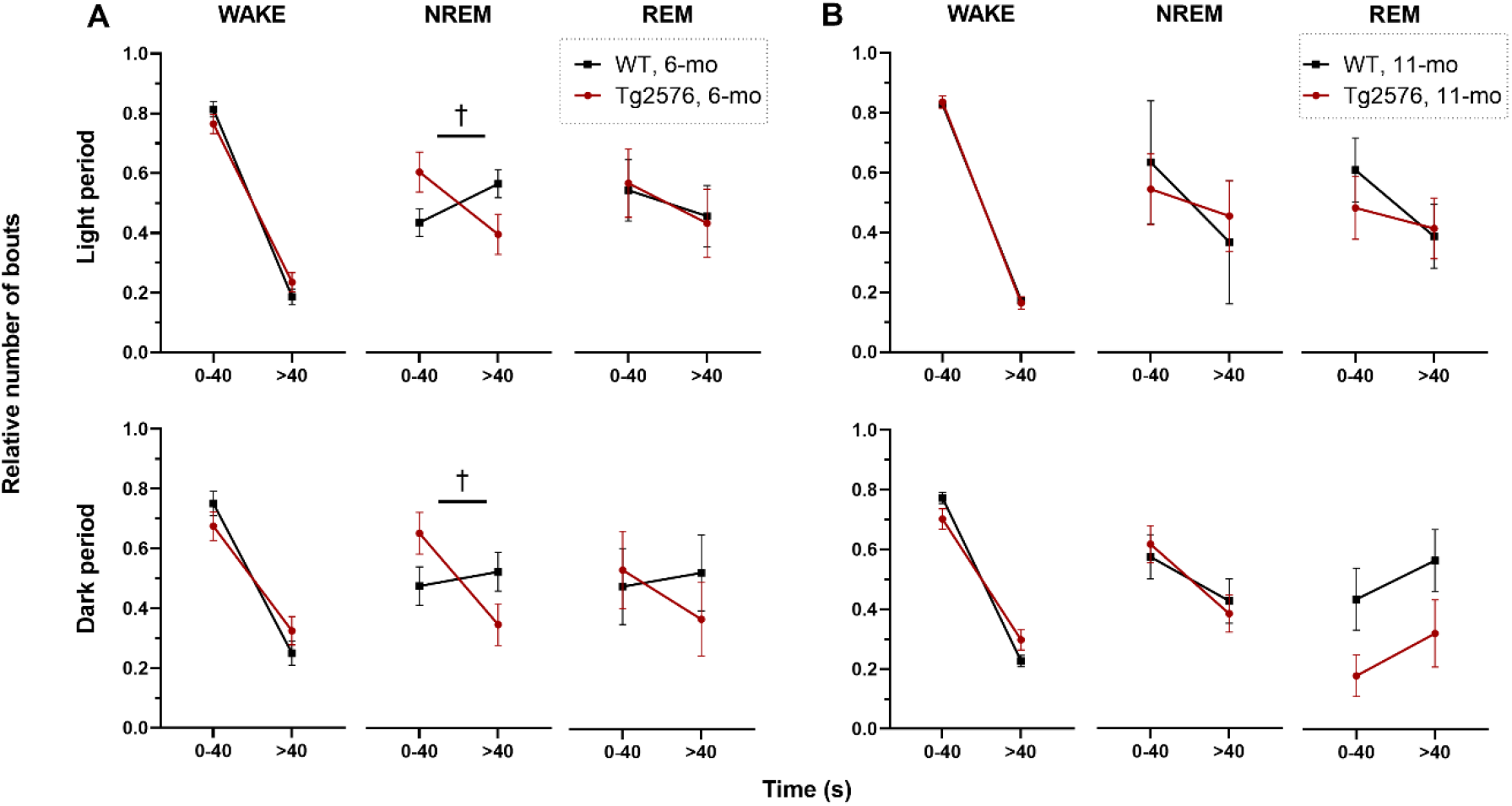
Distribution of bout lengths in all three vigilance states. The length of WAKE, NREM sleep, and REM sleep bouts in light and dark periods from Tg2576 and WT mice were calculated and categorized as short (0-40 s) and long (>40 s) in **A)** 6-mo old and **B)** 11-mo mice. All data are expressed as mean ± SE-, † *p* ≤ .01, mixed ANOVA.

### 3.3. Power spectrum analyses detected age-related differences in Tg2576 mice

Altered spectral power, particularly in theta and alpha waves,^23, 24^ arises as a signature trait in AD.^25^ Thus, we evaluated EEG power spectrum in WAKE, NREM, and REM in 6- and 11-mo Tg2576 and WT mice **(Figure 4)**. We found a significant main effect of genotype on beta band during WAKE **(Table 1)** and pairwise comparisons indicated that the beta power was higher in 11-mo Tg2576 mice compared to their age-matched WT littermates (Tg2576: *M* = 24.55, *SD* = 5.14; WT: *M* = 20.79, *SD* = 4.33, *p* = .036).

**Figure 4.**
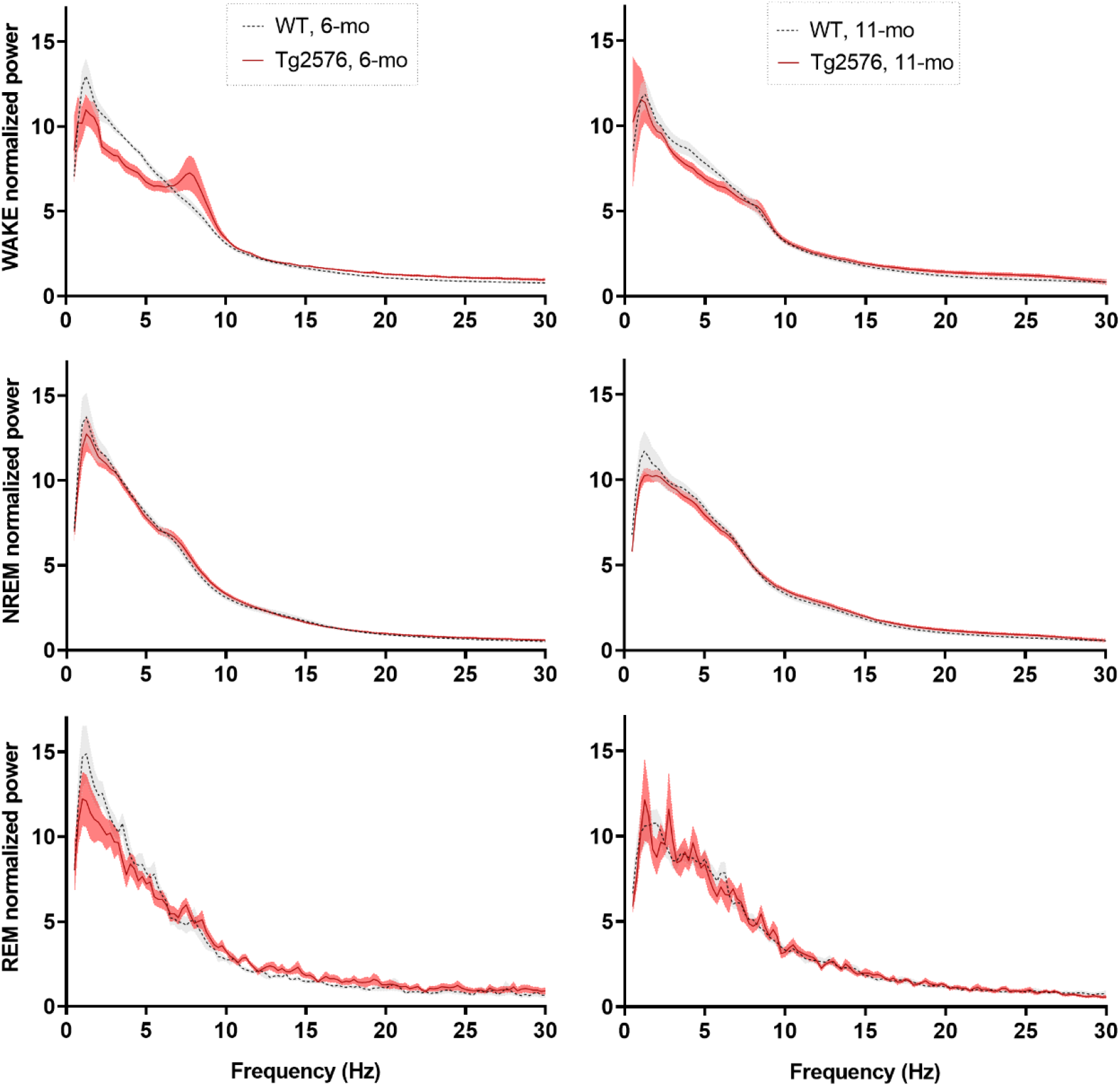
EEG spectral power in WAKE, NREM and REM stages. The data was normalized indicating the percentage of each bin with reference to the total EEG spectral power between 0.5 – 30 Hz in each vigilance state and plotted separately for 6- and 11-mo Tg2576 and WT mice.

**Table 1.** Effect of genotype and age on frequency bands in each vigilance state.

|  |  | WAKE |  |  | NREM |  |  | REM |  |  |
| --- | --- | --- | --- | --- | --- | --- | --- | --- | --- | --- |
| | | *F* (1, 36) | *p* | $\eta_p^2$ | *F* (1, 36) | *p* | $\eta_p^2$ | *F* (1, 36) | *p* | $\eta_p^2$ |
| Delta | Genotype | 3.43 | .072 | .09 | 2.27 | .140 | .06 | 3.04 | .090 | .08 |
|  | Age | 0.19 | .666 | .01 | 5.16 | **.029** | .13 | 5.07 | **.031** | .12 |
| Theta | Genotype | 1.32 | .258 | .04 | <0.01 | .959 | <.01 | 0.02 | .879 | <.01 |
|  | Age | 1.16 | .290 | .03 | 0.50 | .484 | .01 | 6.57 | **.015** | .15 |
| Alpha | Genotype | 3.47 | .071 | .09 | 2.81 | .103 | .07 | 4.25 | **.047** | .11 |
|  | Age | 0.27 | .606 | .01 | 2.56 | .119 | .07 | 7.30 | **.010** | .17 |
| Sigma | Genotype | 1.91 | .176 | .05 | 0.36 | .554 | .01 | 3.18 | .083 | .08 |
|  | Age | 3.06 | .089 | .08 | 4.49 | **.041** | .11 | 5.67 | **.023** | .14 |
| Beta | Genotype | 7.80 | **.008** | .18 | 3.00 | .092 | .08 | 1.31 | .260 | .04 |
|  | Age | 2.75 | .106 | .07 | 4.18 | **.048** | .10 | 0.03 | .855 | <.01 |

*Note:* p values in bold represent a significant result. Statistics were performed by two-way ANCOVA. Frequency bands: delta (0.5–4 Hz), theta (4–8 Hz), alpha (8–12 Hz), sigma (12–16 Hz), beta (16–30 Hz). WAKE = wakefulness; NREM = non-rapid eye movement sleep; REM = rapid eye movement sleep.

Further analysis demonstrated that age, but not genotype, affected the spectral power in both NREM and REM sleep **(Table 1)**. Analysis in NREM spectrum showed a significant main effect of age on delta and sigma bands. Pairwise comparisons revealed that 6-mo Tg2576 mice had significantly higher delta power than 11-mo ones (6-mo: *M* = 159.51 *SD* = 19.43; 11-mo: *M* = 137.71, *SD* = 12.97, *p* = .047), while their sigma power was lower than in older transgenic mice (6-mo: *M* = 29.97, *SD* = 3.43; 11-mo: *M* = 35.98, *SD* = 4.65, *p* = .032). The aging effect on REM was seen as a decrease in delta, and an increase in theta, alpha and sigma power regardless of genotype. Theta power was higher in the older Tg2576 mice (6-mo: *M* = 101.02, *SD* = 13.78; 11-mo: *M* = 113.34, *SD* = 12.50, *p* = .042), and alpha (6-mo: *M* = 46.98, *SD* = 8.03; 11-mo: *M* = 55.25, *SD* = 9.35, *p* = .045) and sigma (6-mo: *M* = 26.73, *SD* = 4.14; 11-mo: *M* = 33.88, *SD* = 7.21, *p* = .023) powers were larger in older WT mice, compared to younger mice within the genotype.

### 3.4. Delta power is altered in Tg2576 mice in a NREM sleep amount-dependent manner

Sleep depth, for which power in delta frequency range (0.5 - 4 Hz) is a thoroughly validated proxy,^26^ has been indicated as a highly predictive measure of the clearance rate in the rodent and human brain.^27, 28^ Therefore, we compared the summed NREM delta power between Tg2576 and WT mice in both age groups in light and dark periods separately.

Our results indicated a lower summed delta power (**Figure 5**) in 6-mo Tg2576 mice in both light (95% CI [51.039, 2703.694], *p* = .042) and dark (95% CI [508.456, 1792.152], *p* = .001) periods compared to age-matched WT controls, whereas in 11-mo mice, summed delta power in Tg2576 was lower only in the dark period (95% CI [29.683, 1286.352], *p* = .041).

**Figure 5.**
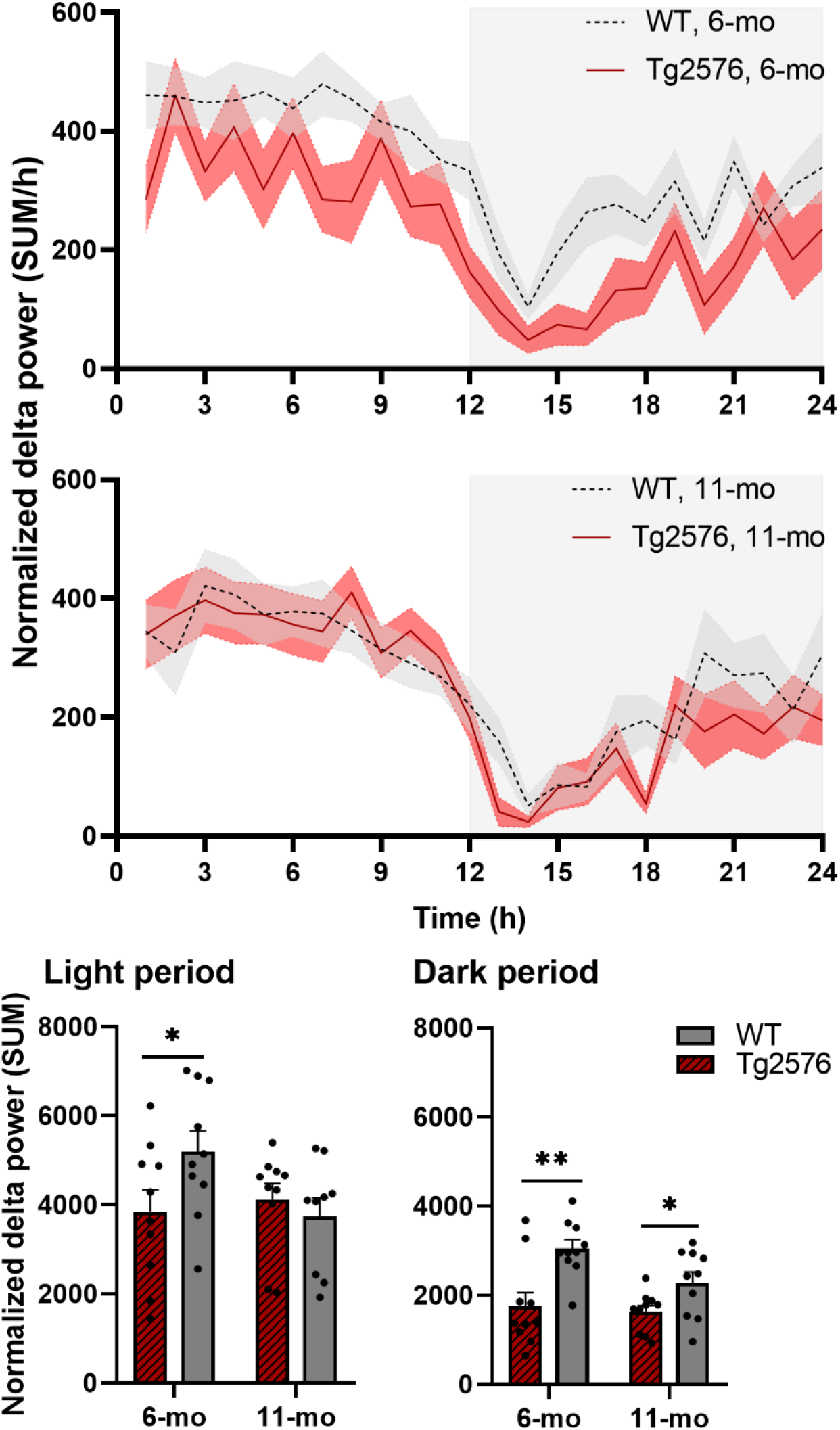
Light and dark delta power patterns in 6- and 11-mo Tg2576 and WT mice. **A)** Sum of delta power during the light and dark periods displayed a significant difference between 6-mo Tg2576 and WT mice (*top panel*). In 11-mo mice (*middle panel*), this difference between genotypes was only present during the dark period (*bottom panel*). All data are expressed as mean ± SE-, *p < .05, **p < .01, two-way mixed ANOVA.

To further examine whether there was a delta power alteration regardless of the time spent in NREM sleep, we additionally analyzed the delta power density per genotype and age in both light and dark periods. We observed no significant differences between the density scores of Tg2576 and their WT controls (**Figure 6**).

**Figure 6.**
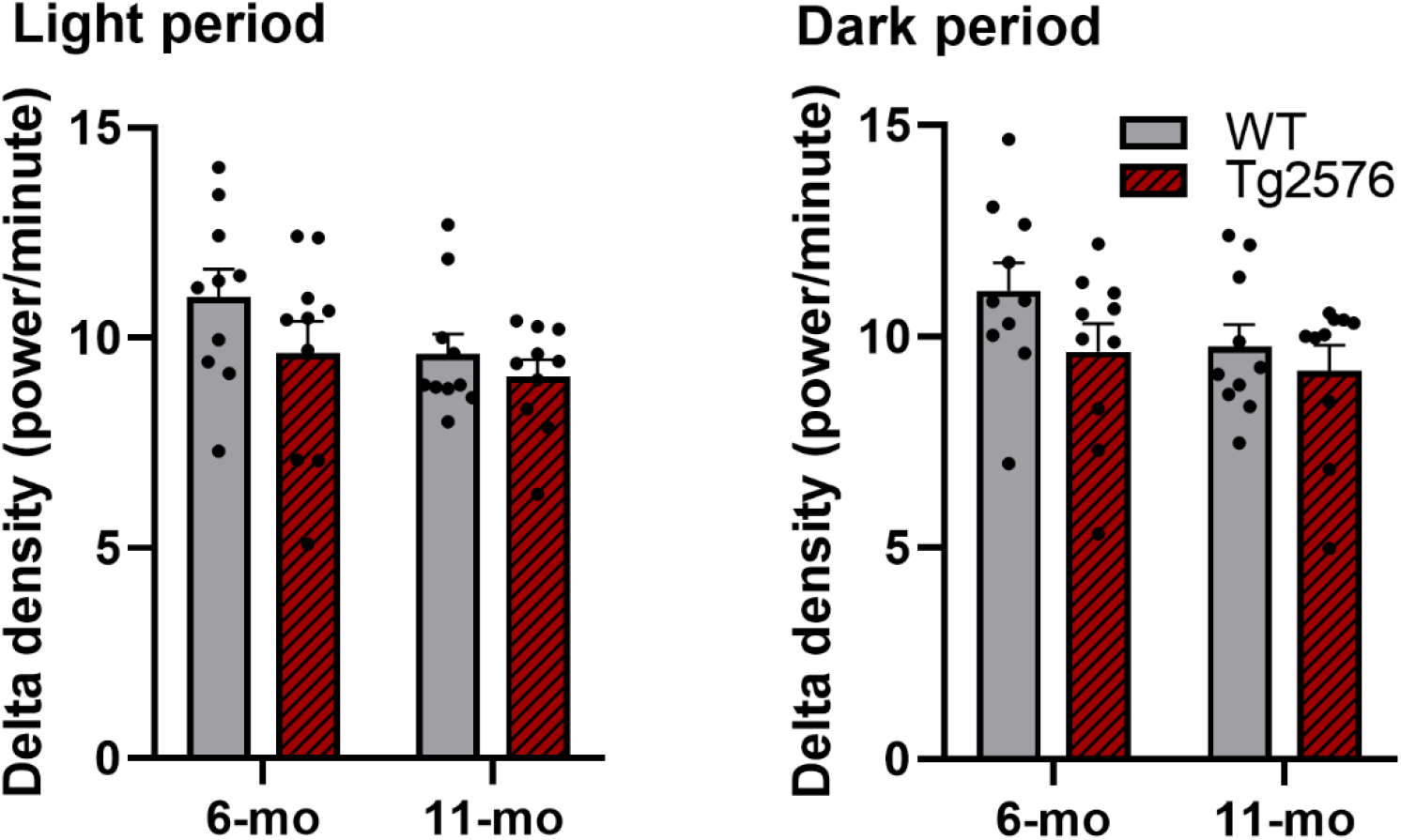
Delta density relative to time spent in NREM. No significant difference at both ages was observed between Tg2576 and WT mice. Data is expressed as mean ± SE-, two-way ANCOVA.

### 3.5. Slow wave energy impairment already appears in plaque-free stage in Tg2576 mice

We further analyzed the homeostatic regulation of SWS by computing the cumulative SWE across 24 h recordings.^29^ Our analysis demonstrated that the intensity of sleep was different in 6-mo Tg2576 mice compared to their age-matched WT controls (*F*(1.300,23.394) = 8.869, *p* = .004, η_p_^2^ = .330, **Figure 7A**) per 24 h. However, such difference did not persist between genotypes in 11-mo mice (**Figure 7B**). We also evaluated the effect of age within each genotype and, although 6-mo WT mice had higher SWE compared to 11-mo animals (*F*(1.228,22.104) = 5.278, *p* = .025, η_p_^2^ = .227, **Figure 7C**), there was no effect of age in Tg2576 mice (**Figure 7D**).

**Figure 7.**
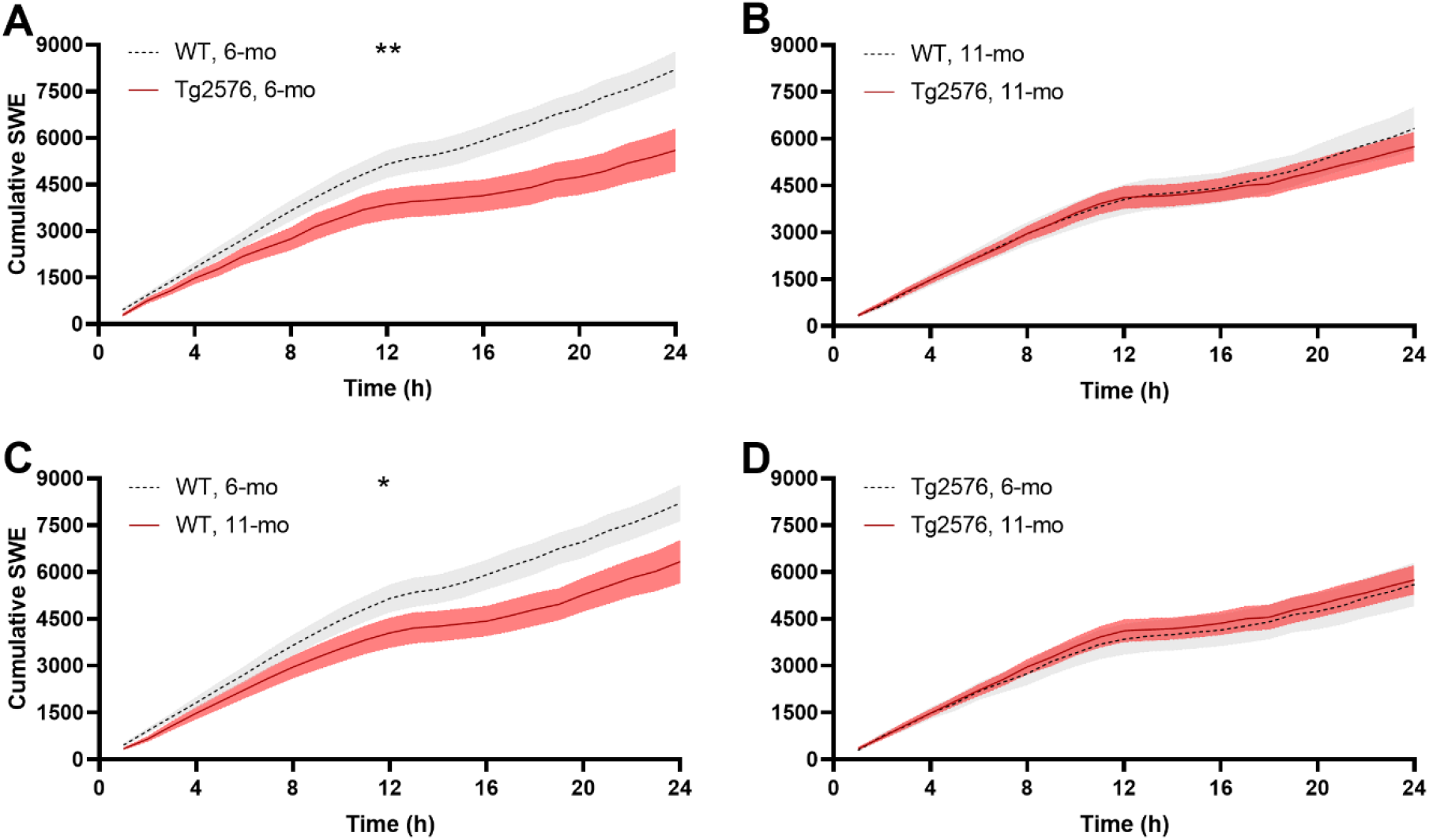
Effects of genotype and age on cumulative SWE. Cumulative SWE differ during the dark period between genotypes in **A)** 6-mo mice, while there is no difference between genotypes in **B)** 11-mo mice. Analysis within the same genotype revealed that SWE decreased in **C**) 11-mo WT mice compared to 6-mo animals. However, **D)** SWE in Tg2576 mice did not differ between the two ages. Comparisons were carried out by a two-way mixed ANOVA (genotype*time or age*time). SWE: slow wave energy.

### 3.6. Plaque burden was confirmed during moderate stage

We then calculated the plaque burden in the hippocampus of each mouse from the sagittal sections stained with Congo red. Our analyses demonstrated that Tg2576 mice in moderate stage showed greater Aβ plaque deposition in comparison to age matched WT animals (*t*(7) = 3.850, *p* = .006; **Figure 8B**), but not in pre-morbid stage (t(6) = 0.169, p = .871; **Figure 8A**).

**Figure 8.**
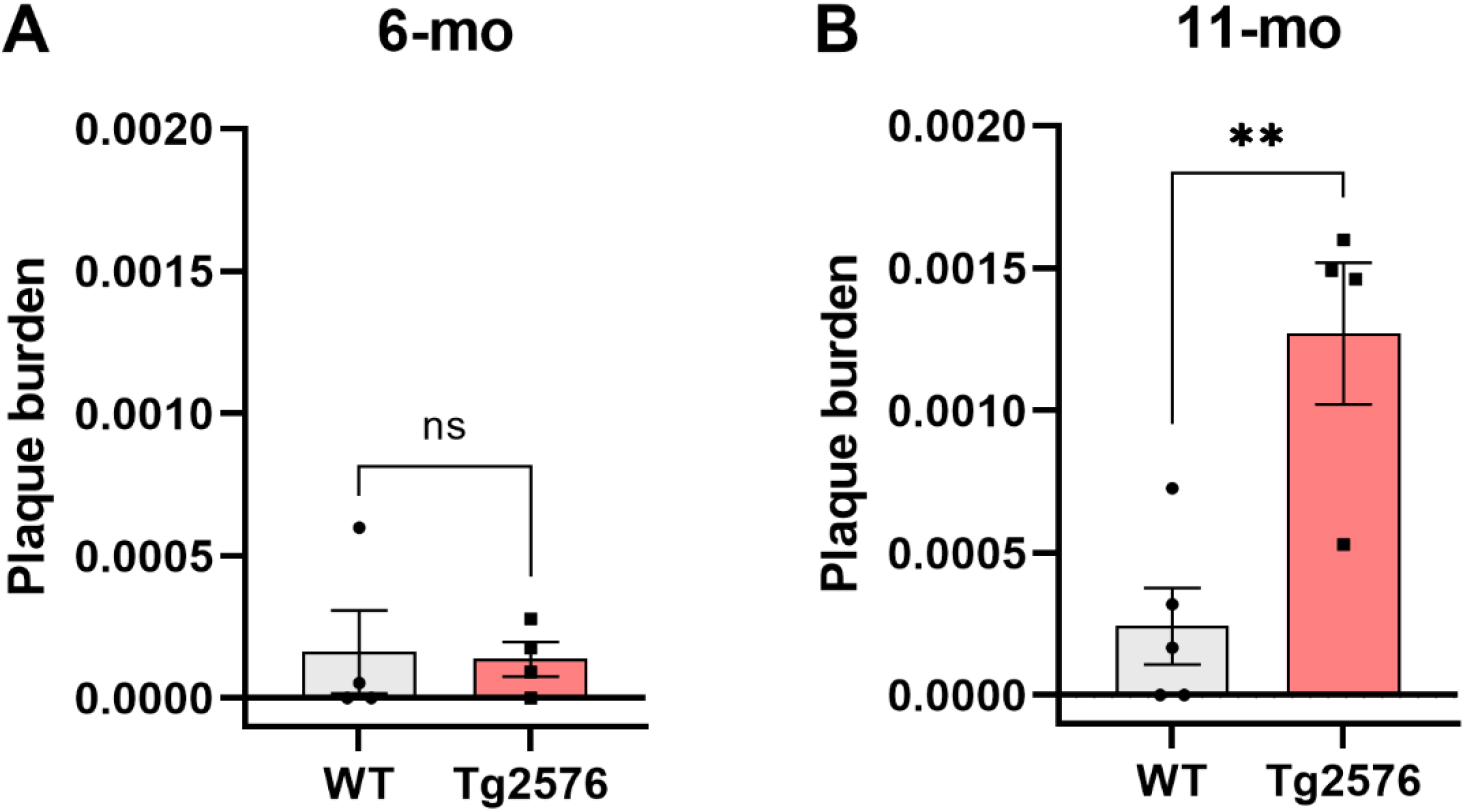
Plaque burden in 6-mo and 11-mo mice assessed by Congo Red staining. A) There was no significant difference between 6-mo Tg2576 and WT mice in plaque burden in the hippocampus. B) Our analysis indicated a significant plaque burden in 11-mo Tg2576 mice compared to WT mice. Data is expressed as mean ± SE-, independent t-test.

## 4. Discussion

Sleep abnormalities are prevalent in AD and other neurodegenerative dementias^30, 31^ with symptoms including insomnia, circadian disorders, excessive daytime sleepiness,^32^ and sleep fragmentation.^33^ Already at early stages of disease, AD patients have decreased SWS, experience longer awakenings and more pronounced sleep fragmentation.^1^ In this study, we report sleep-wake disturbances in the frequently used Tg2576 mouse model of AD, exhibiting many of the age-related disease characteristics observed in humans. Interestingly, we did not find a difference in sleep-wake proportions between Tg2576 and WT mice during the light period (i.e., the resting period in rodents). However, our findings indicate increased wakefulness and decreased NREM sleep during the active dark period in Tg2576 mice already at early age (plaque-free pre-morbid stage) as well as in older ones (plaque-burdened stage) compared to age-matched WT controls. This finding is similar to that of the TgCRND8 AD model mouse,^34^ which is characterized by early plaque formation at 3 months of age and dense-cored plaques by 5 months.^35^ One of the key findings of our study is that during the total (24 h) recording, sleep-wake proportion alterations in Tg2576 were prominent at 6-mo but not at 11-mo, compared to WT mice. The reason for disappearance of genotype-related differences in the 24 h sleep-wake proportions in the aged animals appears to be due to the natural aging phenotype of WT mice. These findings may indicate premature onset of age-related sleep alterations in Tg2576 mice and explain why previous studies in 12-mo Tg2576 animals could not find differences in time spent in each state compared to WT controls.^15^

One of the major effects of aging on sleep-wake patterns is to limit the ability to sustain NREM sleep.^36^ Relative to age-matched WT mice, we detected a trend (*p* = .054) towards higher amounts of short (0 - 40 s) and lower amounts of long (>40 s) NREM bouts in 6-mo Tg2576 mice, indicating an altered sustainability of NREM sleep. Considering the large effect size of this trend, our study was likely underpowered and therefore bigger animal cohorts would likely be required to unveil a significant statistical difference. However, based on these preliminary results, we speculate that the destabilization of NREM sleep in Tg2576 mice may already be present at an early age and may herald a premature age-related sleep phenotype. Moreover, our SWE findings support the notion of an early aging sleep phenotype in Tg2576. Specifically, the cumulative SWE was lower in the 6-mo Tg2576 mice than in the 6-mo WT mice. However, there were no differences between the 11-mo Tg2576 and WT mice. These results indicate that at pre-morbid, plaque-free stage, sleep quality in Tg2576 mice is already altered.

A plausible explanation for the observed sleep-wake alterations present at youth in Tg2576 is the altered dynamics of soluble amyloid beta (Aβ), which has been proposed to affect neuronal networks at wide range^37^ by targeting synapses and disrupting excitatory and inhibitory neurotransmission.^38^ Thereby, our and previous studies’ observations raise the question whether specific sleep-wake alterations, early-onset SWE deficiency and/or decreased sustainability of NREM episodes, could become clinically valuable biomarkers of pre-morbid AD in humans.

Unlike patients with advanced AD,^39–41^ our aged Tg2576 mice showed a decline in delta power in NREM power spectrum at 11-mo compared to younger transgenic mice. It is conceivable that the age groups (6-mo; plaque free and 11-mo; moderate) included in this study do not represent the sleep characteristics of patients in advanced stages of AD. In fact, others observed a trend towards higher delta power in 17-mo Tg2576 mice^13^, suggesting a “slowing” in the EEG spectrum similar to that of AD patients in the advanced stages of the disease.

We examined delta power profiles of Tg2576 and reported a reduced total delta power in both light and dark periods in 6-mo Tg2576 mice compared to 6-mo WT mice. At 11-mo, the total delta power was reduced only in the dark period in the Tg2576 mice, relative to WT controls. Delta power density results, however, pointed out that total delta power is highly dependent on the amount of NREM sleep. Altogether, our data suggest that although power spectrum is similar in terms of average delta frequency between the genotypes, the total delta power of Tg2576 mice is reduced due to their decreased amount of NREM sleep. Therefore, our findings on early-onset altered delta activity may have important implications over protein handling and accumulation in AD. In fact, Aβ levels have been reported to be higher during the dark period in mice, and an interrelation between Aβ and sleep quality reported in both humans and animal models.^11, 42–44^ Thus, a bidirectional relationship between sleep-wake disturbances and AD has been postulated.^45^ This bidirectional relationship between decreased sleep quality and toxic protein accumulation, possibly exacerbating one another, may be one of the grounds for the progression of the disease. Therefore, the direct impact of early age alterations in delta activity on the efficiency of protein handling in vivo, warrants further exploration.

Furthermore, our analysis revealed a greater plaque burden in the hippocampus of 11-mo Tg2576 in comparison to age-matched WT mice. However, 6-mo Tg2576 did not possess amyloid plaques that can be quantified by Congo red staining. These results together with our analyses of sleep indicate that similar to AD patients^46^, sleep-wake problems start before the accumulation of the plaques in Tg2576 mice.

A methodological limitation of this study is the staging accuracy of REM sleep, which is highly dependent on the EMG signal quality. Consequently, abnormalities in the EMG signal caused high variability in the REM sleep scoring in our animals. Another limitation of our report is the lack of purely chronobiologic assessments (i.e. circadian rhythms), hence, we cannot rule out circadian misalignments in this model of murine AD. Additionally, the exploration of early disarrays of sleep-wake patterns in AD models could provide useful information to the field of pre-symptomatic biomarkers of AD.

## 5. Conclusion

Our findings indicate that Tg2576 mice have similar sleep-wake disturbances to AD patients, and that those onset in early, pre-morbid stages of the disease. Our research confirms that animal studies will undoubtedly remain crucial for exploration of sleep-mediated mechanisms in neurodegeneration and provides fruitful information to design future studies considering the age-related sleep differences between AD model and healthy mice. Finally, our findings highlight the importance of designing preclinical and clinical research focusing on improving sleep quality from early ages to potentially reduce the prevalence of symptomatic AD.

## Acknowledgments

This work was supported by the Neuroscience Center Zurich (ZNZ) through the patronage of Rahn and Bodmer Co. (DN) and the Synapsis Foundation for Alzheimer’s Research via an earmarked donation of the Armin & Jeannine Kurz Stiftung (DN). The authors thank Aakriti Sethi for her assistance during sleep recordings, and Ami Beuret for his technical support.

## Author Contributions

SK wrote the initial manuscript; collected the data, performed the statistical analyses, and interpreted the data. CGB and ID contributed to the data acquisition; reviewed and edited the manuscript. DB, DM, JMB, and CRB reviewed and edited the manuscript. DN conceptualized the study; reviewed and edited the manuscript.

All authors approved the final draft.

## Data availability statement

The data that support the findings of this study are available from the corresponding author upon reasonable request.

